# Case study: Genomic characteristics of the gut microbiome, *Campylobacter* and *Salmonella* genotypes in three cases of gastroenteritis co-infections

**DOI:** 10.1101/2025.04.29.651233

**Authors:** Bilal Djeghout, Steven Rudder, Thanh Le Viet, Ngozi Elumogo, Gemma C. Langridge, Nicol Janecko

**Affiliations:** Quadram Institute Bioscience, Norwich Research Park, Norwich, NR4 7UQ, United Kingdom; Centre for Microbial Interactions, Norwich Research Park, Norwich, NR4 7UG, United Kingdom; Eastern Pathology Alliance, Norfolk and Norwich University Hospital, Norwich, NR4 7UY, United Kingdom

**Keywords:** Dysbiosis, microbiome, *Campylobacter*, *Salmonella*, gastroenteritis

## Abstract

*Campylobacter jejuni* and *Salmonella enterica* co-infections in gastroenteritis cases are underexplored, particularly in relation to microbiome dysbiosis. This study uses metagenomic and culture-based approaches to investigate pathogen genotypes and stool microbiome composition. Stool samples from three patients were analysed using culture, qPCR and whole genome sequencing (WGS) of genomes and metagenomes. Strain typing and antimicrobial resistance (AMR) profiling were conducted for both pathogens. The stool microbiome structure was compared between the three patients and across three time points in one patient.

*C. jejuni* and *Salmonella enterica* were of different strains and different proportions in the cases. *C. jejuni* sequence types were identified as ST-794, ST-50 and ST-10025, while *S. enterica* serovars Enteritidis ST-183 and *S.* Derby ST-39 were identified. *C. jejuni* AMR genotypes included *bla*_OXA-447_, *gyr*A (T86I), and *tet*(O). *S.* Enteritidis contained AMR gene *mdsa*A-B, and *S.* Derby contained *mdsa*A-B and *fos*A7 genes.

Despite similarities in Bristol scales for stool composition, the three cases exhibited distinct microbiome population structures, suggesting there is no one signature microbial profile of microbiomes with a *Campylobacter*-*Salmonella* co-infection. A temporal trend towards microbial diversity was observed within a patient, shifting from *Proteobacteria* dominance (93.3%) during acute infection towards *Bacteroidetes* (89.5%) abundance post-infection, suggesting that moving towards diverse communities may play a key role in patient recovery.

These findings reveal a case study of microbiome disruption for patients infected with *C. jejuni* and *S. enterica* and highlight the value of direct stool sequencing in understanding pathogen abundance in stool and microbiome dysbiosis.

## Introduction

Foodborne bacterial infections are a major global health concern, with *Campylobacter* and *Salmonella* contributing as the two leading bacterial pathogens [1]. These infections pose significant public health burdens due to the potential for severe complications, including systemic diseases [2, 3].

*Campylobacter*, predominantly *Campylobacter jejuni*, is the most common bacterial cause of gastroenteritis in the UK [4]. *C. jejuni* is primarily acquired through consuming contaminated food, especially undercooked poultry and poultry products [4]. Clinical manifestations range from mild diarrhoea to more severe outcomes such as Guillain-Barré syndrome and bloodstream infections [4]. Laboratory diagnosis typically involves culture or PCR-based detection, but detection and characterisation challenges arise due to *Campylobacter*’s fastidious growth requirements and its often-low relative abundance in stool samples [5]. Culture-independent methods, such as PCR assays, are commonly used for rapid detection; however, these assays fail to provide comprehensive genotypic information, such as strain subtyping and antimicrobial resistance profiles, important for clinical management and epidemiological surveillance [6, 7].

*Salmonella enterica* are the second leading cause of bacterial gastroenteritis, also primarily associated with the consumption of contaminated food, including eggs, meat, and fresh produce [8, 9]. While most infections are self-limiting, *Salmonella* can cause invasive disease, particularly in vulnerable populations [8–10]. Like *Campylobacter*, diagnostic approaches for *Salmonella* typically involve a combination of rapid PCR assays and culture. While culture methods remain the gold standard, PCR-based diagnostics are commonly used due to their speed and sensitivity. However, these methods are limited in tracking antimicrobial resistance and providing detailed strain-level data [7, 11–13].

Co-infections involving *Campylobacter* and *Salmonella* are not often reported, and the clinical significance of these events remains unclear [14, 15]. Experimental evidence from murine models shows that co-infections with *C. jejuni* and *S.* Typhimurium can exacerbate the symptoms of gastroenteritis and have an increased bacterial load of *C. jejuni* while *S.* Typhimurium levels remain stable [14]. Co-infection with *C. jejuni* and entero-invasive *Escherichia coli* (EIEC) was shown to promote *C. jejuni* colonisation; however, the effect diminished over time [14]. A study investigating these co-infection dynamics using human and avian epithelial cell lines found that the presence of *E. coli* K12 negatively impacted the metabolic activity of *C. jejuni* strain 11168, highlighting the complex interactions between bacteria during co-infection [16].

Disruptions in the gut microbiome are closely linked to gastrointestinal infections, as they can significantly change the composition of the lower gut microbial community [17]. During infection, beneficial populations such as *Firmicutes* tend to decrease, while opportunistic pathogens like *Proteobacteria* can increase [18]. This disruption can lead to reduced microbial diversity, which has been observed in independent *Campylobacter* and *Salmonella* infections [19, 20]. Both pathogens have been shown to alter the gut microbiome, though the specifics of these changes remain under investigation [18].

Identifying campylobacteriosis and salmonellosis requires laboratory examination of stool samples alongside clinical assessment. Diagnostic laboratory testing involves a multi-pathogen rapid PCR assay followed by a reflexive isolate culture and sequencing; however, for *Campylobacter,* diagnostic laboratories may not conduct culture due to resources, thereby limiting full characterisation of this pathogen [11, 12]. While culture-independent testing offers rapid and precise detection results at the genus or species level, PCR-based methods do not provide comprehensive genotypic information such as sub-typing and antimicrobial resistance genotype profiles [7, 21, 22]. This may limit effective management of patient treatment [23, 24]. Exclusively using culture-independent testing also limits the ability to accurately track both established and emerging *Campylobacter* and *Salmonella* strains, essential for effective public health surveillance.

In recent years, metagenomic testing and analytical tools have become invaluable for characterising the gut microbiome and identifying some pathogen genotypes without the need for culture. The method enables thorough analysis of microbial populations within a sample, and specialised bioinformatic protocols can offer insights into the genetic traits of pathogens, including antimicrobial resistance and virulence factors [25–27]. The utility of direct metagenomic sequencing from stool samples demonstrated how clinically relevant *Campylobacter* attributes, including strain subtyping and resistance profiles can be assessed [28].

This case study aimed to: 1) analyse *Campylobacter* and *Salmonella* genotypes responsible for three identified co-infection cases using culture and direct sequencing methods at the acute phase of infection, and assess the utility of methods in detecting, identifying, and characterising the pathogens; 2) compare bacterial stool microbiomes between three gastroenteritis cases caused by *Salmonella*-*Campylobacter* co-infections, identifying shared or distinct bacterial patterns and 3) characterise temporal disruption in bacterial diversity within one patient’s microbiome over time.

## Materials and methods

Ethical approval for this study was obtained from the University of East Anglia Research Ethics Committee [Ref 2018/19-159]. Research involving human tissue (stool) complied with Norwich Biorepository license NRES number – 19/EE/0089, under the IRAS Project ID - 259062 approved by the UK Human Tissue Authority (HTA). The National Health Service (NHS) Eastern Pathology Alliance (EPA) network diagnostic laboratory in Norwich, United Kingdom (UK) provided excess diagnostic stool samples for this project from the Norfolk and Waveney, UK catchment area. As part of routine screening for infectious intestinal diseases, the EPA laboratory used a rapid automated PCR-based culture-independent testing panel (Gastro Panel 2, EntericBio Serosep, Crawley, United Kingdom) to test submitted stool samples for various pathogens, including *Campylobacter* (*C. jejuni, C. coli and C. lari)* and *Salmonella* according to their standard operating procedures. The inclusion criteria for this study were concurrent positive PCR results for *Campylobacter* and *Salmonella* within the same stool sample. Three cases matching the inclusion criteria were identified and followed up as case studies between August 2022 and May 2023.

### Stool sample collection

Three cases were included in this study. Patient-1, a 57-year-old female, presented in the emergency department but was not admitted as an inpatient. The initial sample was collected on August 18, 2022. The research nurse practitioner conducted a survey, and supplementary samples were submitted at two weeks and 15 weeks post-symptom onset (Figure S1).

Patient-2, a 68-year-old male and patient-3, a 27-year-old male submitted stool samples to on the 16^th^ and 17^th^ of May 2023, respectively, after a general practitioner office visit. No further samples were collected post-infection recovery for patient-2 and patient-3 (Figure S1).

Once PCR results were available at the EPA diagnostic laboratory, excess stool samples from each patient were transported to the Quadram Institute Bioscience research laboratory for microbiological isolation of *Campylobacter* and *Salmonella,* metagenome DNA extraction and sample preservation (Figure S2).

### i) Isolation and short-read sequencing of genomes

***Campylobacter* isolation:** Each stool sample underwent *Campylobacter* isolation by direct culture using a modified ISO method (EN ISO 10272 - 2019) for detecting and enumerating *Campylobacter* [29]. In brief, a 10 μl of stool was directly plated to modified charcoal-cefoperazone deoxycholate agar (mCCDA), supplemented with cefoperazone and amphotericin-B supplements (Oxoid, Hampshire, UK). To maintain a microaerophilic environment, all plates involved in the isolation protocol were placed in anaerobic jars with CampyGen 2.5L sachet (Oxoid, Hampshire, United Kingdom) and incubated at 37°C for 48 hours. *C. jejuni* strain 81116 [30] was used as a positive control throughout the protocol. Following incubation, up to 20 suspected *Campylobacter* colonies per sample were transferred to a second mCCDA plate for purification and further sub-cultured onto Columbia blood agar supplemented with 5% horse blood (Oxoid, Hampshire, UK). Presumptive *Campylobacter* isolates were identified based on typical colony morphology and oxidase testing (Thermo Fisher Scientific, Loughborough, UK).

***Salmonella* isolation:** *Salmonella* was isolated from each stool sample by direct plating. In brief, 10 μl of stool was plated to each side of a bi-plate containing Brilliance™ *Salmonella* Agar and Xylose Lysine Deoxycholate (XLD) agar (Oxoid, Hampshire, UK). Plates were incubated at 37°C for 24 hours. Suspected *Salmonella* colonies per agar type were sub-cultured onto MacConkey agar (Oxoid, Hampshire, UK) and incubated at 37°C for 24 hours. Each suspected *Salmonella* colony was purified on Tryptic Soy Agar (TSA) Tryptic Soy Agar (Oxoid, Hampshire, UK) and incubated at 37°C for 24 hours. Presumptive *Salmonella* isolates were identified based on typical colony morphology throughout the protocol.

**Genome DNA extraction and sequencing:** Presumptive *Campylobacter* and *Salmonella* isolates underwent DNA extraction using Maxwell RSC Cultured Cells DNA Kits (Promega, Hampshire, UK), following the manufacturer’s instructions. DNA libraries were prepared using the Illumina DNA Prep kit (Illumina Ltd, Cambridge, UK), as detailed in a previous study [31], and paired-end (PE150) libraries were sequenced on an Illumina NextSeq500 instrument with a mid-output flowcell (NSQ® 500 Mid output KT (v2) (300 CYS) (Illumina, Cambridge, United Kingdom).

***Campylobacter* and *Salmonella* genome analysis:** Raw sequence reads were stored on an in-house instance of IRIDA [32]. Trimming of reads was performed with fastp (v0.19.5) [33]. Genus and species prediction for *Campylobacter* used Kraken2 (v2.1.1) [32]. Typing of *Salmonella* genomes was conducted with SeqSero2 [34]. Assembly of paired-end reads of all genomes was conducted using SPAdes (v3.12.0) [35], and the quality of assemblies was assessed through QUAST (v5.0.2) [36]. Multi-locus sequence typing (MLST) was carried out using the PubMLST database (v2.16.1) [37] to determine the sequence type (ST) based on genome assemblies. abriTAMR AMR gene detection pipeline with AMRFinderPlus was used to detect AMR determinants [38]. SNP calling was performed using Snippy (v3.2) and Snippy-core (v3.2) (https://github.com/tseemann/snippy), with the highest-quality isolate randomly selected from one patient serving as the reference genome for *Campylobacter* and *Salmonella*, respectively. RAxML (v8.2.4) [39] was used to construct a phylogenetic tree based on SNPs identified in the core genomes of *Campylobacter* and *Salmonella* isolates.

### ii) Sample preparation and metagenome DNA extraction

**Raw stool preparation:** An adapted host depletion method was used to minimise the amount of human DNA present as previously described [40, 41]. Briefly, the digestion of host DNA was completed by adding 200 µl of buffer (comprising 5.5 M NaCl and 100 mM MgCl2 in nuclease-free water) to 200 µl of stool, followed by adding 35 µl of 1% saponin (Tokyo Chemical Industry UK Ltd, Birkenhead, United Kingdom) and 10 µl of HL-SAN DNase (Articzymes, Tromsø, Norway). Samples were thoroughly mixed and incubated at 37 °C with agitation at 6010 G for 20 minutes.

Upon completion of the digestion process, samples underwent a wash step using 300 µl phosphate-buffered saline (containing NaCl [58.44 g/mol], KCl [74.55 g/mol], Na_2_HPO_4_ [141.96 g/mol], and KH_2_PO_4_ [136.09 g/mol]) and then centrifuged at 18,900 G for 5 minutes. The supernatant was carefully removed. The resulting pellet was designated as the input stool sample and underwent complete DNA extraction.

The Maxwell® RSC Fecal Microbiome DNA Kit (Promega, Hampshire, UK) was used for the metagenome DNA extraction following the manufacturer’s instructions. Subsequent DNA eluent was quantified using Qubit fluorometer with the dsDNA quantitation high sensitivity kit (Life Technologies Ltd, Paisley, UK).

### iii) Campylobacter and Salmonella quantification

***Campylobacter* quantification:** A quantitative PCR (qPCR) assay was conducted on each metagenome DNA sample using the LightCycler® 480 Probes Master kit on the LightCycler® 480 Instrument II (software LCS480 v1.5.0.39) (Roche Diagnostics Ltd, West Sussex, UK). The *CadF* target gene was selected for *Campylobacter* quantification [42, 43]. In brief, each reaction comprised a 20 μl reaction mixture containing 2 μl of DNA sample, 10 μl of a primer-probe master mix, 7 μl of nuclease-free water, 0.4 μl of *cadF* forward and reverse primers (10 μM each), and 0.2 μl of cadF probe (10 μM) [42]. qPCR cycling conditions included a pre-amplification step at 95°C for 10 minutes to initialize the reaction, an amplification phase of 45 cycles, with each cycle involving denaturation at 95°C for 15 seconds, followed by annealing and extension at 55°C for 1 minute and a cooling step at 44°C for 30 seconds to facilitate post-reaction stabilisation. Samples with cycle threshold (CT) values in a range of 1 to 40 cycles were considered positive for *Campylobacter*.

***Salmonella* quantification:** *Salmonella* quantification in stool metagenome DNA samples was conducted using the PMA Real-Time PCR Bacterial Viability Kit – *Salmonella enterica* (*inv*A) (Biotium, Inc, Fremont CA, USA) on the LightCycler® 480 Instrument II (software LCS480 v1.5.0.39) (Roche Diagnostics Ltd, West Sussex, UK).

Each reaction consisted of a 20 μl reaction mixture containing 2 μl of the DNA sample, 10 μl of Fast EvaGreen Master Mix, 6 μl of nuclease-free water, and 2 μl of *inv*A forward and reverse primers (5 μM each) [44]. qPCR cycling conditions included a pre-incubation step at 95°C for 5 minutes to initiate the reaction, amplification phase of 40 cycles, with each cycle involving denaturation at 95°C for 5 seconds, followed by annealing and extension at 60°C for 30 seconds. A melt phase consisted of 1 cycle with 3 steps: denaturation at 99°C for 1 second, annealing at 57°C for 1 second, and a final denaturation at 99°C. Following the amplification and melt phases, a cooling step was performed at 37°C for 1 second. Samples with CT values between 1 to 40 were considered positive for *Salmonella*.

### iv) Metagenome DNA sequencing and analysis

**Short-read metagenome DNA library preparation and sequencing:** An adapted metagenome DNA library preparation method utilised the Illumina DNA Prep (M) tagmentation kit in accordance with the manufacturer’s instructions (Illumina Inc., San Diego CA, USA). Further quality control (QC) procedures were implemented on DNA libraries. This included sample quantification with Qubit™ 1X dsDNA High Sensitivity (HS) using Qubit fluorometry (Winsford, United Kingdom) to determine DNA concentration and the assessments of insert size and molar concentration to ensure the integrity and suitability of the libraries for sequencing.

Paired-end indexed libraries were sequenced on the Illumina NovaSeq PE150 platform by Novogene Ltd., Cambridge, UK, to achieve a minimum sequencing depth of 8 GB per sample.

**Abundance analysis of *Campylobacter* and *Salmonella* in stool microbiomes:** Raw sequence reads were stored on the in-house instance of IRIDA [32]. Initial quality control was performed was performed of the sequencing reads using the QIB Clean-up pipeline v1.4 (https://github.com/telatin/cleanup). Potential human read contamination was further removed using Hostile (v1.1.0) [45] against the human-t2t-hla reference database, and quality filtered using TrimGalore (v0.6.10) [46]. The complete pipeline used for the human decontamination and QC is available at: https://github.com/aponsero/QIB_WGScleanup. Raw reads were then subjected to quality control and adapter trimming using fastp (v0.19.5) [33]. MetaPhlAn4 (v4) [47] was employed for taxonomic profiling and involved mapping pre-processed reads to the MetaPhlAn4 marker gene database, enabling the determination of relative abundances of bacterial taxa present in the samples. The abundance of the microbiome was visualised using R Studio [48] _ using the ggplot2 (v3.4.0) package for generating bar plots and the dplyr (v1.1.3) package for data manipulation.

**Metagenome-derived genome (MDG) of *Campylobacter* and *Salmonella* analysis:** *Campylobacter* and *Salmonella* MDGs were assembled following a previously described method [41]. Briefly, reads classified at the genus level were extracted from Kraken2 (v2.1.1) [49] and Bracken (v2.6.0) [50] outputs, treated as isolate reads, and assembled using Shovill (v1.1.0) (https://github.com/tseemann/shovill). The quality and completeness of the MDGs were evaluated using BUSCO (v3.0.2) [51] and CheckM (v1.0.11) [37], with genome characteristics assessed using QUAST (v5.0.2) [52]. Sequence types were determined by multi-locus sequence typing (MLST, v2.16.1) with the PubMLST database [53], and antimicrobial resistance determinants were analysed using AMRFinderPlus (v3.11.4) [54].

**Alpha and beta diversity:** Alpha and beta diversity analysis was conducted using Phyloseq (v1.46.0) [55] and vegan (v2.6-4) packages in R 4.3.0. Alpha diversity metrics such as species richness, Shannon diversity index, and Simpson diversity index were calculated to assess within-sample diversity [56]. Beta diversity analysis calculated the differences between samples based on the microbial composition [57]. Diversity was visualised using Principal Component Analysis (PCA) and plotted with PC1 and PC2 axes.

## Results

Stool samples from three patients [patient-1; patient-2; patient-3]were analysed. All three patients submitted samples during the acute gastroenteritis symptom stage identified as time point 1 (T1), and patient-1 submitted two further samples [at post-recovery of gastroenteritis (T2) and 15 weeks post symptoms (T3)] (Table 1).

**Table 1.**
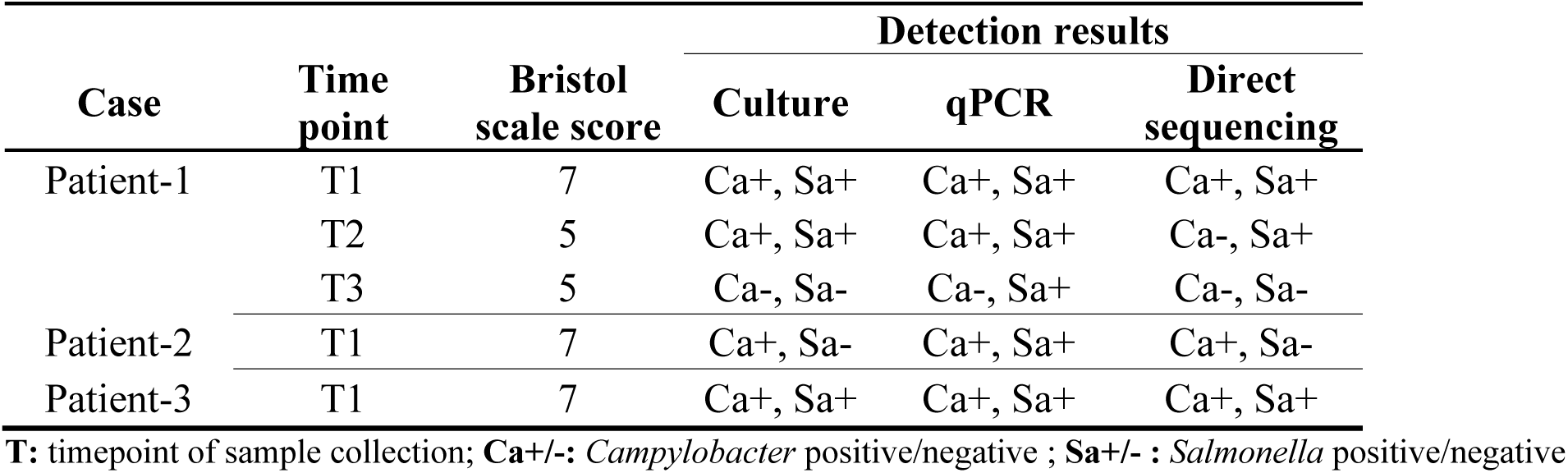
Summary of *Campylobacter* and *Salmonella* detection by culture, qPCR, and direct sequencing in three co-infection case study patients.

| Case | Time point | Bristol scale score | Detection results |  |  |
| --- | --- | --- | --- | --- | --- |
|  |  |  | Culture | qPCR | Direct sequencing |
| Patient-1 | T1 | 7 | Ca+, Sa+ | Ca+, Sa+ | Ca+, Sa+ |
|  | T2 | 5 | Ca+, Sa+ | Ca+, Sa+ | Ca-, Sa+ |
|  | T3 | 5 | Ca-, Sa- | Ca-, Sa+ | Ca-, Sa- |
| Patient-2 | T1 | 7 | Ca+, Sa- | Ca+, Sa+ | Ca+, Sa- |
| Patient-3 | T1 | 7 | Ca+, Sa+ | Ca+, Sa+ | Ca+, Sa+ |

**T:** timepoint of sample collection; **Ca+/-:** *Campylobacter* positive/negative ; **Sa+/- :** *Salmonella* positive/negative

At T1, all patients experienced symptoms consistent with acute gastroenteritis with diarrhoeal stool as the main sign, sought medical attention and subsequently submitted stools classified as Bristol scale score 7. Subsequent samples of patient-1 were classified as Bristol scale score 5. Culture testing obtained *Campylobacter* and *Salmonella* isolates in patients-1 and −3, while for patient-2 only *Campylobacter* isolates were recovered. qPCR assays identified both pathogens in all patient samples at the symptomatic stage of the infection (Table 1).

### Genotype characterisation of Campylobacter and Salmonella

***Campylobacter* spp.:** A total of 48 *C. jejuni* isolates [patient-1: n=24; patient-2: n=18; patient-3: n=6] were recovered by direct culture (Figure 1). Each patient harboured a different *Campylobacter* sequence type (ST). *Campylobacter* DNA quantity in patient-1 remained the same at T1 and T2 with a real-time PCR Ct value of 30 (∼10⁶ CFU/mL). No detectable amount of *Campylobacter* was identified at T3. In patient-2, the Ct value was 36 (∼10⁴ CFU/mL); in patient-3, the Ct value was 40 (∼10² CFU/mL).

**Figure 1.**
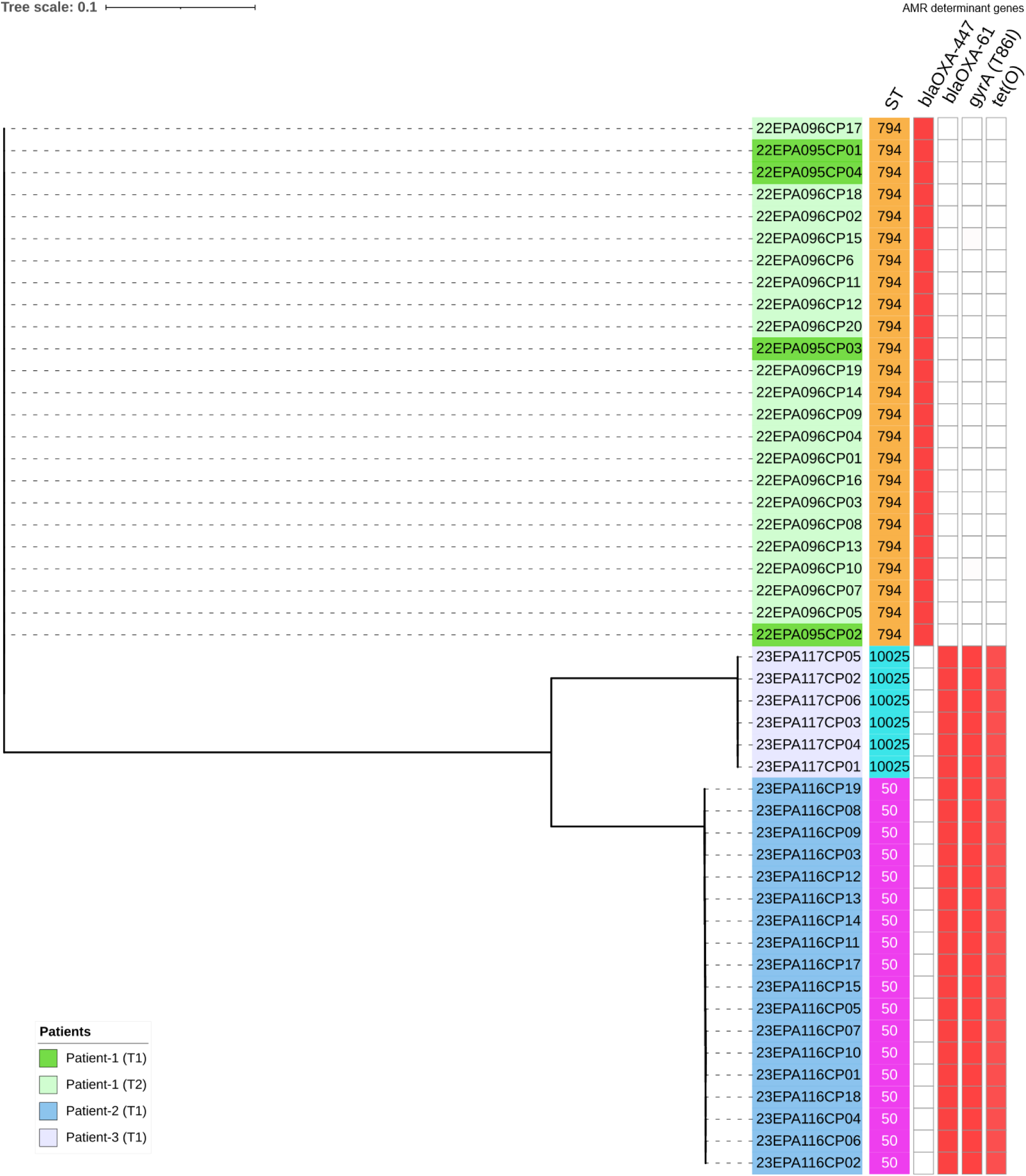
A maximum likelihood core genome SNP-based phylogenetic tree of 48 *C. jejuni* isolates from three patients co-infected with *Campylobacter* and *Salmonella*. Isolate IDs are coloured by patient ID and time point. ST is indicated, and the presence of AMR gene determinants is indicated by red blocks.

In the stool sample of patient-1 at T1 (acute gastroenteritis, PCR+), four *C. jejuni* ST-794 isolates were recovered, all containing the AMR gene *bla*_OXA-447_. Similarly, at T2 (12 days post initial symptoms, PCR+), 20 *C. jejuni* ST-794 isolates were obtained, all carrying the AMR gene *bla*_OXA-447_. No *Campylobacter* was isolated at T3 (15 weeks post initial symptoms, PCR-).

The stool sample of patient-2 at T1 (acute gastroenteritis, PCR+) contained 19 *C. jejuni* ST-50 isolates, all containing AMR genotype profile with *bla*_OXA-193_, *tet*(O), and *ars*P genes and the *gyr*A (T86I) mutation (Figure 1).

For patient-3 at T1 (acute gastroenteritis, PCR+), six *C. jejuni* ST-10025 isolates were recovered, all harbouring AMR genes *bla*_OXA-193_, *tet*(O), *ars*P, *sat*4, and *aph(3’)-IIIa* and the *gyr*A (T86I) mutation. Core genome single nucleotide polymorphism (SNP) analysis of *C. jejuni* isolates revealed distinct inter-patient genomic patterns, while intra-patient genomes were highly related (Figure 1). In patient-1, one *C. jejuni* ST-794 isolate from timepoint T2 exhibited a single SNP difference. In patient-2, *C. jejuni* ST-50 isolates had SNP differences that ranged from 1-6 SNPs. *C. jejuni* ST-10025 from patient-3 isolates exhibited SNP differences that ranged from 1-5 SNPs.

***Salmonella* spp.:** A total of 18 *Salmonella* isolates were recovered by direct culture [patient-1: n=9 (*S.* Enteritidis); patient-2: n = 0; patient-3: n = 9 (*S.* Derby)].

In patient-1, eight isolates were identified as *S.* Enteritidis ST-183 [three from T1 and five from T2] and one isolate from T2 was an unspecified ST. No isolates were recovered from patient-1 at T3. In patient-3, nine isolates were identified as *S.* Derby ST-39 at T1 only.

Quantification of *Salmonella* DNA in the stool of patient-1 at T1 was a Ct value of 31.23 (∼10⁵ CFU/mL), 22.11 (∼10⁵ CFU/mL) at T2 and 35 (∼10³ CFU/mL) at T3. Quantification of *Salmonella* DNA for patient-2 was a Ct value of 35 (∼10³ CFU/mL) at T1 and 29.71 (∼10⁷ CFU/mL) for patient-3 at T1.

All *S.* Enteritidis genomes exhibited the AMR genes *mdsA* and *mdsB* at T1 and T2 (Figure 2) and exhibited SNP differences ranging from 46 to 54 at T1. Genomes from T2 showed SNP differences ranging from 0 to 5 among the six isolates. Intra-patient SNP difference between T1 and T2 ranged from 22 to 37 SNPs.

**Figure 2.**
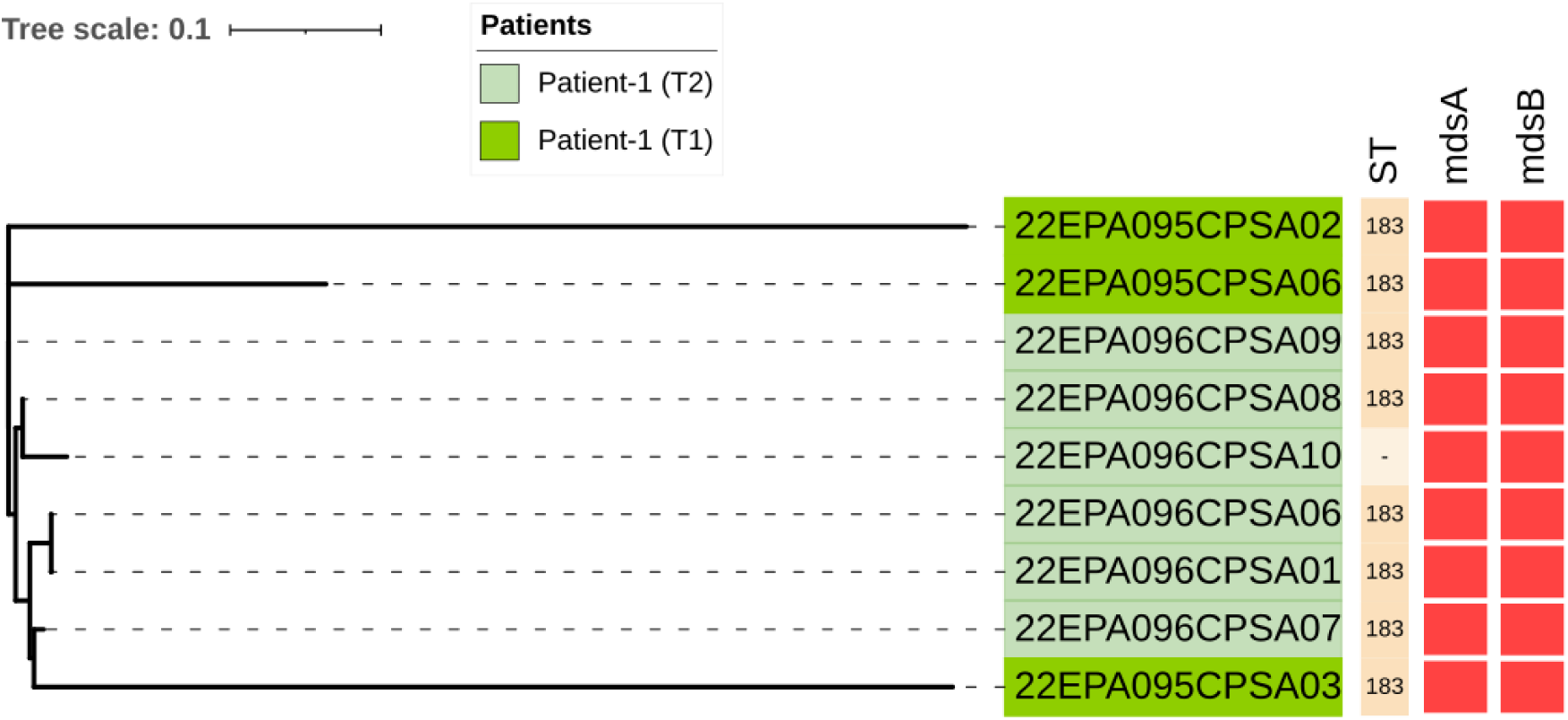
A maximum likelihood core genome SNP-based phylogenetic tree of nine *Salmonella* Enteritidis isolates from a co-infected with *Campylobacter* and *Salmonella*. Isolate IDs are coloured by the patient ID and time point. ST are indicated and the presence of AMR gene determinants is indicated by red blocks.

For patient-3 at T1, SNP differences among the *S.* Derby genomes ranged from 79 to 181 SNPs (Figure 3). All genomes of *S.* Derby contained AMR genes *mdsA*, *mdsB* and *fosA7*.

**Figure 3.**
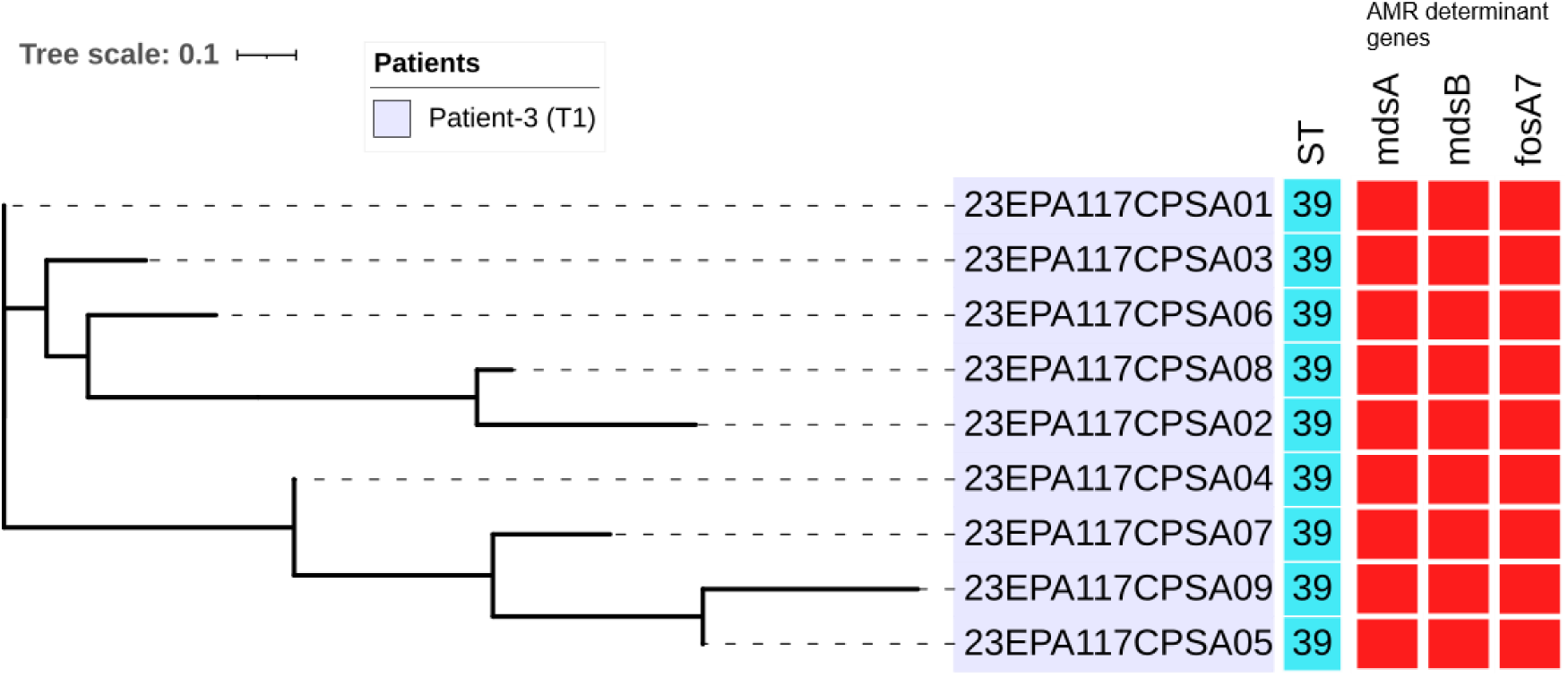
A maximum likelihood core gnome SNP-based phylogenetic tree of *Salmonella* Derby isolates from a co-infected patient with *Campylobacter* and *Salmonella*. Isolate IDs are coloured by the patient ID and time point. ST are indicated and the presence of AMR gene determinants is indicated by red blocks.

## Inter-patient microbial population structure and diversity

Interpatient stool sample comparison at T1 revealed notable differences at the phylum and family level (Figure 4 A and B). In the sample of patient-1, the microbial population proportions consisted of *Firmicutes* (65%), *Proteobacteria* (20%) and *Bacteroidetes* (15%). In contrast, patient-2 and patient-3 samples, the proportion of *Bacteroidetes* was high at 89.07% and 82.82%, respectively. A difference was observed in the proportion of *Proteobacteria* detected in patient 3 (15.33%) compared to patient-2 (<1%), while *Firmicutes* proportions were similar (1.06% and 1.83%, respectively). *Actinobacteria* were detected at a low proportion in patient-2 (0.15%) and patient-3 (0.06%) but were not detected in patient-1 (Table S2).

**Figure 4.**
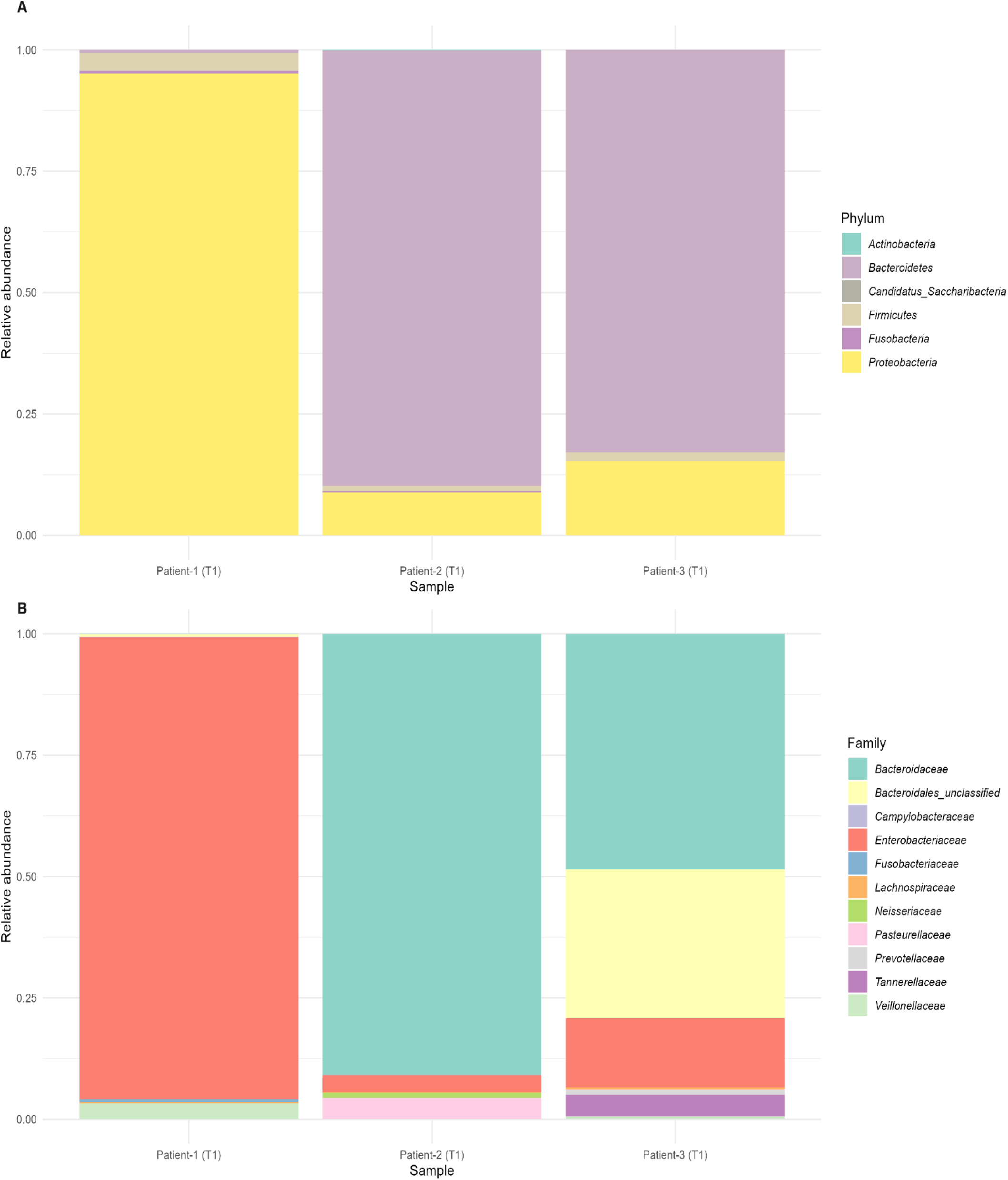
Relative abundance by proportion of microbiome population structure of stool samples for three patients’ co-infection with *Campylobacter* and *Salmonella*. Relative abundance proportions profiles are shown at the **(A)** Phylum level and **(B)** Family level, highlighting the top 10 most abundant families plus *Campylobacteraceae*.

At the family level, the same trends were observed. *Bacteroidaceae* was the most dominant group in patient-2 and patient-3, representing 90% and 77.4% of relative abundance, respectively, whereas the *Bacteroidaceae* proportion was minor (0.65%) in patient-1 (Figure 4B). *Lachnospiraceae* and *Enterobacteriaceae* were proportionally high in patient-1, accounting for 40% and 20%, respectively, yet lower in comparison to patient-2 and to patient-3. *Tannerellaceae* was more abundant in patient-3 (4.2%) compared to patient-1 and patient-2, where it was nearly absent. Other families such as *Neisseriaceae* and *Streptococcaceae* were detected in the patient-2 sample but were absent in patient-1 and patient-3. Similarly, *Prevotellaceae* was found in low abundance in patient-3 but not in the other two patient samples (Table S2).

At the initial time point (T1) (Figure S3), *Campylobacter* genus reads were detected in all patients: patient-1 had 37,026 reads (0.05% of total genus reads), patient-2 had 34,175 reads (0.06% of total genus reads), and patient-3 had 26,894 reads (0.05% of total genus reads) (Figure S4). *Salmonella* genus reads were present in patient-1 T1 sample with 7,637 reads (0.01% of total genus reads), patient-2 had no *Salmonella* reads, and patient-3 had 43,452 reads (0.08%). In patient-1, *E. coli* was derived from 71,126,933 reads, representing the dominant family *Enterobacteriaceae*, which comprised 95% of the total bacterial reads.

The stool microbiomes of the three patients at T1 exhibited varying levels of *Campylobacter* and *Salmonella* (Table S3). At T1, *Campylobacter* was detected in all three patients, whereas *Salmonella* was only detected by rapid PCR in patient-2 (Table S3). The completeness of *Campylobacter* MDGs ranged from 2.20 to 46.3%, while *Salmonella* MDG was only obtained to 4.3% in patient-3. Additionally, *E. coli* MDGs were recovered from patient-1 and patient-3 (Table S3).

Alpha diversity analysis at T1 revealed distinct differences in microbial diversity across patients (Figure 5). Patient-1 exhibited the lowest diversity, with Shannon and Simpson Diversity Index values of 0.479 and 0.191, respectively. Patient-2 showed slightly higher diversity, with Shannon and Simpson Diversity Index values of 0.879 and 0.328. Patient-3 had the highest diversity among the three, with Shannon and Simpson Diversity Index values of 2.223 and 0.832 (Figure 5). Beta diversity analysis revealed clear separation of microbial communities across the patients at T1 (Figure 6) and demonstrated distinct clustering of microbial profiles for each patient

**Figure 5.**
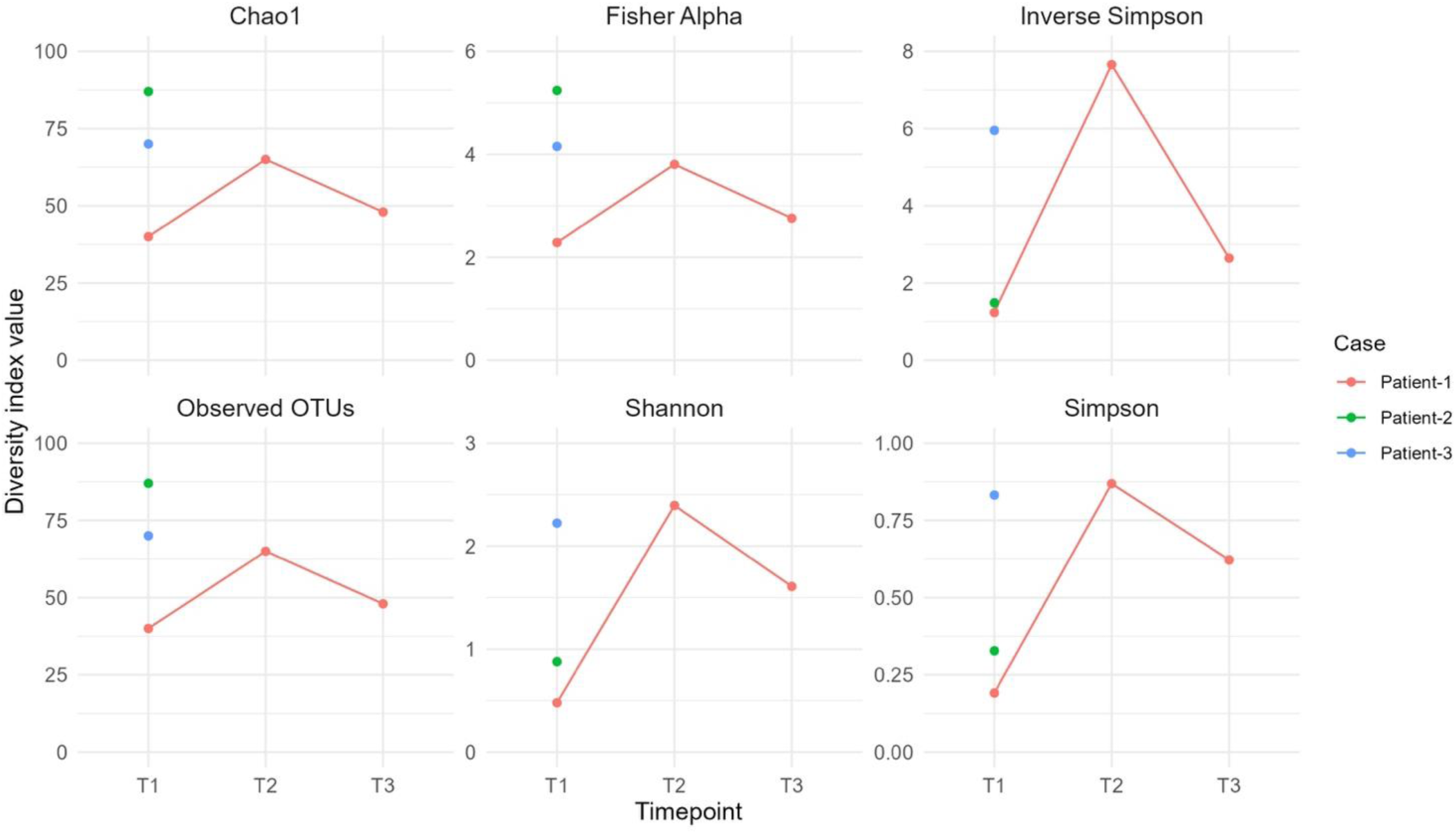
Alpha diversity indices of the stool microbiome over timepoints for three patients during a *Campylobacter* and *Salmonella* co-infection. The figure displays temporal changes in Chao1 (estimated species richness), Fisher Alpha (diversity index), Inverse Simpson (evenness), Observed OTUs (number of unique taxa), Shannon (diversity and evenness), and Simpson (dominance) indices at timepoints T1, T2, and T3 in patient-1 (red dot and line). Patients-2 (green dot) and patient-3 (blue dot) at T1.

**Figure 6.**
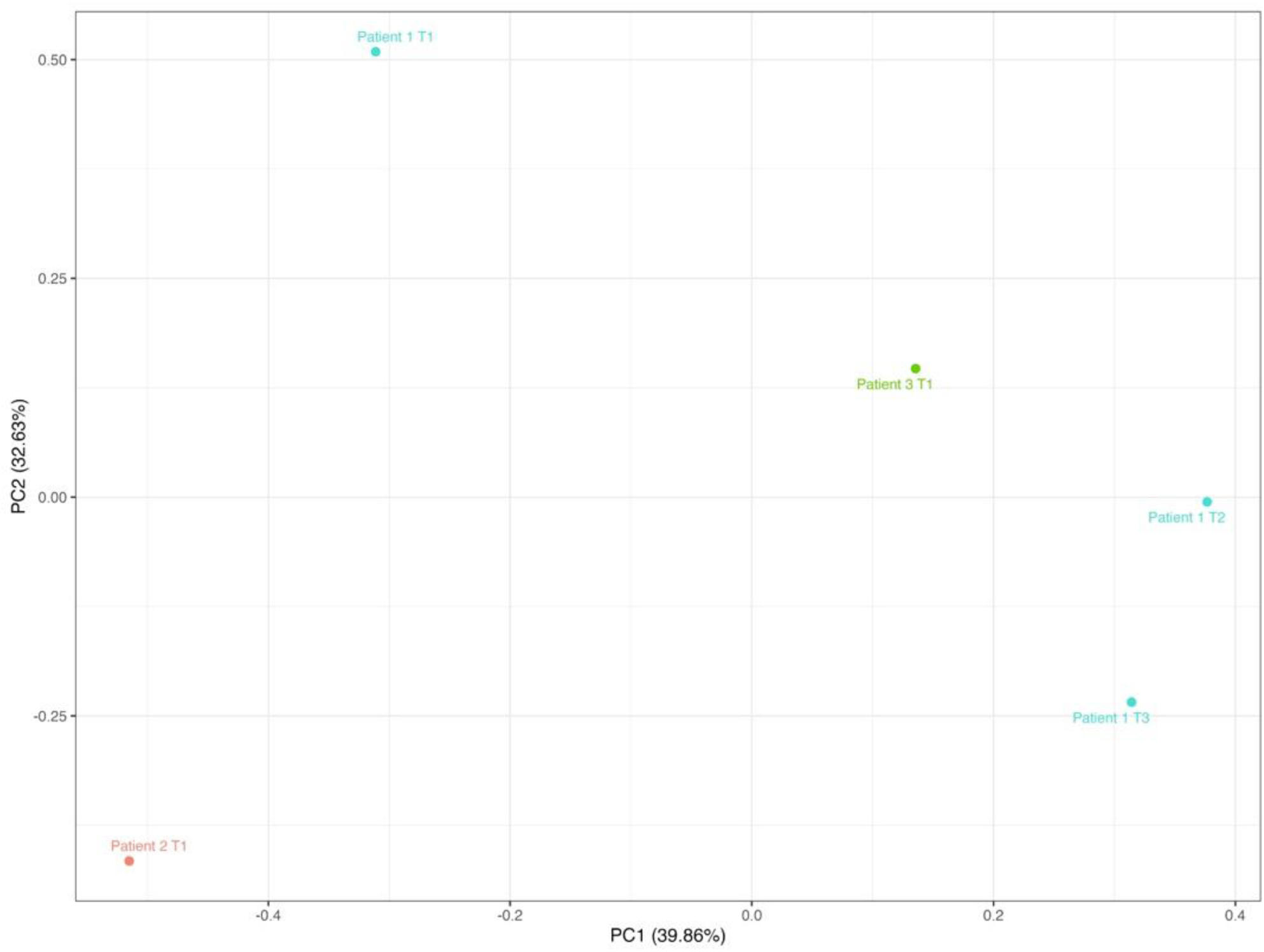
Beta diversity analysis of stool microbiomes in three patients: principal component 1 (PC1) versus principal component 2 (PC2). The plot displays the first two principal components (PC1 vs PC2), accounting for 39.86% and 32.63% of the variance, respectively. Points represent individual samples, with colours corresponding to patient-1 (blue), patient-2 (red), and patient-3 (green).

### Stool microbiome population structure

## Temporal analysis of patient-1

Short-read sequencing of the stool microbiome at three time points revealed fluctuations in the relative proportions of each bacterial population between T1 [acute gastroenteritis] and T2 and T3 [post-infection recovery]. At T1, *Proteobacteria* were the most dominant phylum, representing 93.3% of total reads (71,201,621 reads), followed by *Firmicutes* (5.5% of total reads) and *Bacteroidetes* (1.2% of total reads; 495,717 reads). At T2, the population proportions changed to a predominant *Bacteroidetes* abundance (89.5% of total reads; 110,114,421 reads), followed by *Firmicutes* (7.1%; 7,953,241 reads) and *Proteobacteria* (3.4%; 1,628,403 reads). At T3, *Bacteroidetes* continued to be the predominant phyla in the microbiome (88.2%, 56,074,644 reads), while *Firmicutes* and *Proteobacteria* accounted for 5.4% and 3.6% respectively (Figure 7, Table S2).

**Figure 7.**
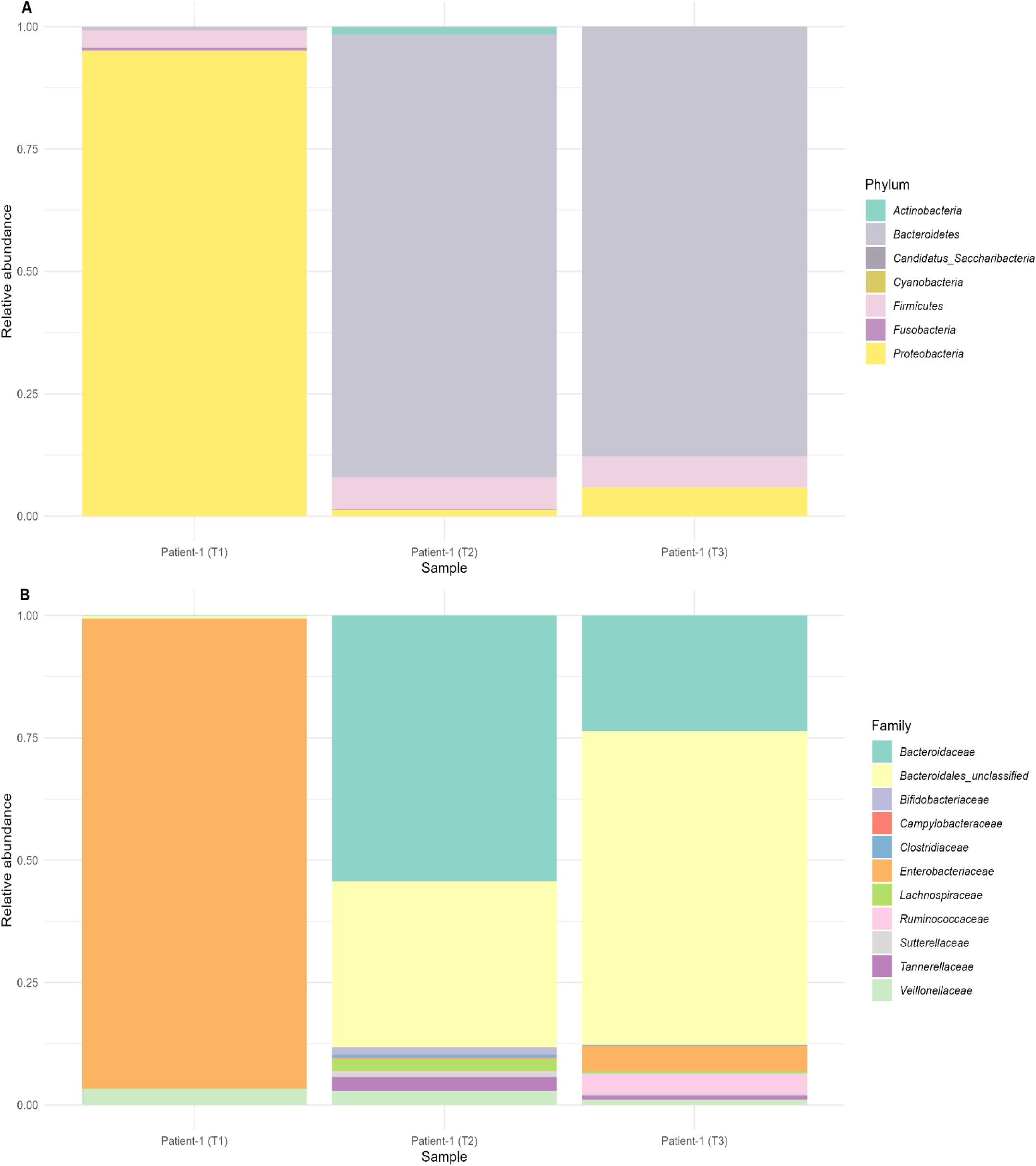
Microbiome relative abundance structure in patient-1 at three timepoints (T1, T2, T3) during and after a *Campylobacter* and *Salmonella* co-infection. Relative abundance profiles are shown at the **(A)** Phylum level and **(B)** Family level, highlighting the top 10 most abundant families plus *Campylobacteraceae*.

At the family and genus taxonomic levels, 95% of bacterial reads were classified as belonging to the *Enterobacteriaceae* family at T1, with the genus *Escherichia* predominating. *Salmonella* accounted for 7,637 (approximately 0.01%) of the total *Enterobacteriaceae* reads. No *Salmonella* MDG was recovered, but the species *Salmonella enterica* was identified from the reads. The relative abundance of *Salmonella* was lower at T2 and T3. Reads from the *Bacteroidaceae* family accounted for 0.65% at T1, increasing to 88% and 87% at T2 and T3, respectively.

At T1, 0.05% (37,026 reads) were assigned to the genus *Campylobacter*. A *Campylobacter* MDG was reconstructed to a 46.3% genome completeness and classified as *C. jejuni*. No further characterisation of ST or resistance profiles was possible. At T2, no MDGs were recovered for *Campylobacter* and *Salmonella*, however *C. jejuni* and *Salmonella enterica* were still identified in the taxonomic classification by MetaPhlAn4. Since *Enterobacteriacea* dominated the microbiome at T1, analysis of *E. coli* was additionally conducted. *E. coli* O15H18, ST-69 MDG was reconstructed with 96% completeness and represented 0.3% of the relative abundance (Table S3). At T3, no MDGs were recovered for *Campylobacter*, *Salmonella*, or *E. coli*.

## Discussion

Our case study targeted the observational comparison of bacterial population composition and relative abundance in the stool of three patients in the context of *Campylobacter* and *Salmonella* co-infections. We identified three distinct *Campylobacter* strains (*C. jejuni* ST-794, ST-50, and ST-10025) and two different *Salmonella* serovars (*S. Enteritidis* ST-183 and *S. Derby* ST-39) across the patients, with varying abundances in the microbiome. Antimicrobial resistance determinants to antimicrobials used for gastrointestinal infections, namely fluoroquinolones were detected in the *Campylobacter* genomes [58]. Furthermore, two out of three patients contained AMR genotypes that conferred resistance to three different antimicrobial classes. Notably, we identified *Campylobacter jejuni* ST-50, ranked among the top ten sequence types across Europe and the UK, underscoring its clinical relevance [59, 60]. The presence of *Salmonella* Enteritidis, a top-ranked *Salmonella* in human infections in the UK, alongside *S.* Derby, further emphasises the clinical significance of these co-infections [61, 62]. While we could not determine the specific etiological contributions of these strains in this study, future research could shed light on whether strain-specific traits play a role in shaping microbiome disruption during co-infection. Likewise, further information would be required on the antimicrobial use and clinical management of patients to understand the clinical impacts of the varied antimicrobial resistance class determinants found in our case study cases. Antimicrobials, to our knowledge, were not prescribed to the case study patients; however, if antimicrobials were used in co-infection scenarios, the varied antimicrobial resistance determinants in both pathogens found in our study would influence the survival and proliferation of either pathogen, thereby potentially complicating clinical outcomes.

*C. jejuni* was detected in higher proportions of microbiome relative abundance compared to *Salmonella* enterica across all patients at T1, although the relative abundance proportions varied among the three patients. This observation aligns with previous studies that have reported variable pathogen loads in co-infections [63]. The highest levels of both *C. jejuni* and *S. enterica* were observed in patient-1; however, this does not necessarily suggest any biological synergy between the two pathogens. Rather, it likely reflects a co-infection with a higher overall pathogen burden, potentially due to factors such as the initial pathogen ingestion load, timing of sample collection concerning symptom onset and peak pathogen load.

The findings from this case study highlight differences in microbial population structure and diversity of stool microbiomes between patients during *Campylobacter* and *Salmonella* co-infections, and differing trends in composition within a patient over time. While diminishing diversity of the gut microbiome has been widely reported in other gastrointestinal infections, such as *Clostridioides difficile* (*C. difficile*) infections, where gut microbiota dysbiosis is a hallmark due to the disruption caused by *C. difficile* colonisation and toxin production, this study is the first to document *Salmonella* and *Campylobacter* dysbiosis in three cases [64].

The temporal trend of microbial population structure was from low to high diversity from acute gastroenteritis to recovery, which aligned with findings from studies on gut microbiota recovery [65–67]. For instance, research on children recovering from watery diarrhoea showed that while antibiotic treatment initially delayed increases in alpha-diversity, diversity eventually increased significantly during the second week of recovery, approaching levels similar to those in non-antibiotic groups [65]. Similarly, studies on children with rotavirus infections demonstrated a transition from a diseased to a healthy microbiota state over time, with temporal variability being larger in infected children than in healthy ones [67]. Additionally, investigations into enteric infections have found that follow-up samples after recovery exhibited more diverse gut microbiota compared to infection stages, characterized by an increase in Bacteroidetes and Firmicutes phyla [66]. These studies collectively support the trend of increasing microbial diversity during recovery from acute gastroenteritis.

The predominance of *Escherichia* during the acute stages of a co-infection has been documented by others, where a bloom of fast-growing facultative anaerobes mostly *Enterobacteriaceae* overpopulates the microbiome [68–70]. This suggests that a transition in the microbial composition takes place and is influenced by host immune responses, leading to a restructuring of the microbial population [65]. The altered environment may then encourage the growth of facultative anaerobes like *Escherichia* and *Streptococcus*, which can thrive in these conditions and may dominate the early stages of an infection [65]. Although we did not identify this phenomenon in all patients, the *Enterobacteriaceae* dominant proportion (95%) in patient 1 aligns with the observations of other studies that observed a similar surge in *Enterobacteriaceae* in enteric infections [69, 71, 72].

Notably, despite being symptom-free yet still testing positive for the pathogens, patient-1 exhibited an increase in microbial diversity by 15 weeks post-symptom onset, indicating dynamic changes in the microbiome composition. This observation aligns with previous findings on gut microbiomes during recovery from diarrhoea, where the recovery phase typically entailed a gradual augmentation in taxonomic richness and diversity [68, 73, 74].

The transition from *Enterobacteriaceae* to *Bacteroidaceae* over time was striking and aligns with studies on keystone microbial families in the microbiome. This shift suggests a restoration of microbial balance following diarrhoeal pathogenesis [74]. Our findings in the temporal trends of patient-1 aligns with the concept that *Bacteroides* species, as keystone species for microbiome recovery, play a crucial role in restoring gut microbiota after infectious diseases causing diarrhoea [74, 75]. During the recovery phase, *Bacteroides* use host-derived nutrients to establish themselves, facilitating the repopulation of other beneficial commensals and restoring microbial diversity and functionality [74–77].

Despite some similarities in the co-infection cases, all three patients exhibited distinct microbial population structures, suggesting there is no singular signature profile of microbiomes during a *Campylobacter-Salmonella* co-infection. The differences between patient microbiomes may not be fully dependent on the pathogen presence but rather may be due to host attributes. The diversity found among the three cases underscores that further investigation is needed into host factors such as diet, age, demographics, and co-morbidities, which have been previously shown to contribute to microbiome population structure differences [78, 79]. Further research is needed to explore host-microbe-community interactions in larger cohorts to understand the roles of strain types, quantities and host factors. Although our metagenomic analysis provides detailed insights into bacterial diversity and pathogen genotypes, it does not capture other microbial contributors, such as viruses (including bacteriophages) and fungi. These organisms also have important roles in microbial interactions and host immune responses. The exclusion of these organisms therefore limits our ability to characterise the complexities of co-infection dysbiosis [80].

In conclusion, our case study of three patients highlights the complex bacterial differences in stool microbiome population structure and function during *Campylobacter* and *Salmonella* co-infections and demonstrates temporal changes within a patient microbiome from dysbiosis to recovery.

## Data Summary

All supporting data and protocols are included in the article or are available as supplementary data files. The online version of this article contains four supplementary figures (Figures S1– S4) and three supplementary tables (Table S1–S3). All sequenced *Campylobacter* and *Salmonella* isolate data are available in the National Centre for Biotechnology Information (NCBI) Sequence Read Archive under the Bioproject accession number PRJNA1231000. Sequence read archive (SRA) accession numbers and associated metadata can be found in the supplementary material of this study (Table S1).

## List of abbreviations

%: percent
°C: Degree Celsius
AMR: antimicrobial resistance
*bla*_OXA_: beta-lactamase oxacillinase gene
C. jejuni: Campylobacter jejuni
CBA: Columbia blood agar
DNA: deoxyribonucleic acid
EPA: Eastern Pathology Laboratory
*gyrA*: DNA gyrase subunit A
HTA: Health Technology Assessment
ISO: International Organization for Standardization
mCCDA: modified Charcoal Cefoperazone Deoxycholate Agar
ml: millilitre
MDG: metagenome derived genome
MLST: Multilocus sequence typing
PBS: Phosphate-buffered saline
PCR: polymerase chain reaction
S.: enteritidis: Salmonella enteritidis
Spp.: species
ST: sequence type
tet: tetracycline gene
UK: United Kingdom
v: Version
WGS: whole genome sequencing
β-lactam: Beta lactam
μl: microlitre
μm: micrometre

## Declarations

### Ethical Approval

This project was approved by the Faculty of Medicine and Health Sciences Research Ethics Committee of the University of East Anglia (FMH REC reference: 201819-159HT).

## Consent for publication

Not applicable.

## Authors contributions

B.D. and N.J. designed the study conception. B.D., N.J. and G.L. conceptualised the plan for the manuscript. B.D. drafted the manuscript. S. R. contributed to *Salmonella* analysis and interpretation. All authors contributed to the edits of the manuscript. B.D. designed figures. T.L.V. and B.D. analysed data and produced figures on R Studio. B.D. and S.R. carried out bioinformatic genomic data analysis and interpretation.

## Acknowledgement

The author(s) gratefully acknowledge the support of the Biotechnology and Biological Sciences Research Council (BBSRC). We extend our sincere gratitude to the clinical nurse (Carmen Walker, in Memorium), the microbiology lab and the administrative team at the Eastern Pathology Alliance network diagnostic laboratory in Norwich, UK, for providing access to samples, with special thanks to Nuno Pedro for assisting in obtaining metadata. We acknowledge the project scientists, laboratory managers and technicians at Quadram Institute Bioscience (QIB) for their invaluable technical support; the QIB core services sequencing team for their expertise in sequence library preparation, quality control, and sequencing, and the QIB core bioinformatics team for their assistance in uploading the DNA sequencing data to NCBI.

