## Supplementary data for "Case study: Genomic characteristics of the gut microbiome, *Campylobacter* and *Salmonella* genotypes in three cases of gastroenteritis co-infections"

***Patient-1 brief case study survey narrative***

A follow-up brief case study survey was conducted with patient-1. Patient follow-up was not possible for patients-2 and -3.

Patient-1 reported first becoming ill on August $7^{th}$, 2022, experiencing symptoms including diarrhoea with light-coloured loose stool, abdominal pain/cramps, slight blood in stool, body aches, joint pain, fatigue, and loss of appetite. The T1 sample was submitted within this time and classified as Bristol scale 7. Ten days later, the patient’s symptoms improved and returned to work. No recent travel or antibiotic use before the onset of illness was reported at the time of the survey. The patient reported no other gastrointestinal or chronic diseases. A subsequent stool sample was collected once symptoms improved at day 12 (T2) and again 15 weeks after initial symptoms (T3). At the 15-week follow-up survey, the patient expressed that bowel function fluctuated, and the stool remained light-coloured. The stool was classified as Bristol scale score 5.

Regarding food consumed prior to symptoms, an open-ended question of foods consumed within 7-days of symptoms was asked. The patient described consuming a rice dish with chicken within 7 days before becoming ill. However, a full food recall survey was not conducted. The patient reported handling rice with chicken dish that ‘went off’ but did not eat that dish. Animal exposure was to a pet dog; they had no other animal exposure before becoming ill.

**Table S1.** Sequence Read Archive accession numbers for stool metagenomes and genome isolates of *Campylobacter* and *Salmonella* recovered from human gastroenteritis samples.

| Sample Name | BioProject | SRA accession number | Source | Type |
|---|---|---|---|---|
| 22EPA095CP | PRJNA1231000 | SRR32586498 | Human stool | Metagenome |
| 22EPA096CP | PRJNA1231000 | SRR32682504 | Human stool | Metagenome |
| 22EPA115CP | PRJNA1231000 | SRR32682503 | Human stool | Metagenome |
| 23EPA116CP | PRJNA1231000 | SRR32682502 | Human stool | Metagenome |
| 22EPA095CP1 | PRJNA1231000 | SRR32682501 | Human stool | Metagenome |
| 22EPA095CP2 | PRJNA1231000 | SRR32681814 | Human stool | Genome |

| | | | | |
|---|---|---|---|---|
| 22EPA095CP3 | PRJNA1231000 | SRR32681813 | Human stool | Genome |
| 22EPA095CP4 | PRJNA1231000 | SRR32681802 | Human stool | Genome |
| 22EPA096CP1 | PRJNA1231000 | SRR32681791 | Human stool | Genome |
| 22EPA096CP10 | PRJNA1231000 | SRR32681780 | Human stool | Genome |
| 22EPA096CP11 | PRJNA1231000 | SRR32681809 | Human stool | Genome |
| 22EPA096CP12 | PRJNA1231000 | SRR32681808 | Human stool | Genome |
| 22EPA096CP13 | PRJNA1231000 | SRR32681807 | Human stool | Genome |
| 22EPA096CP14 | PRJNA1231000 | SRR32681806 | Human stool | Genome |
| 22EPA096CP15 | PRJNA1231000 | SRR32681805 | Human stool | Genome |
| 22EPA096CP16 | PRJNA1231000 | SRR32681804 | Human stool | Genome |
| 22EPA096CP17 | PRJNA1231000 | SRR32681803 | Human stool | Genome |
| 22EPA096CP18 | PRJNA1231000 | SRR32681801 | Human stool | Genome |
| 22EPA096CP19 | PRJNA1231000 | SRR32681800 | Human stool | Genome |
| 22EPA096CP2 | PRJNA1231000 | SRR32681799 | Human stool | Genome |
| 22EPA096CP20 | PRJNA1231000 | SRR32681769 | Human stool | Genome |
| 22EPA096CP3 | PRJNA1231000 | SRR32681798 | Human stool | Genome |
| 22EPA096CP4 | PRJNA1231000 | SRR32681758 | Human stool | Genome |
| 22EPA096CP5 | PRJNA1231000 | SRR32681754 | Human stool | Genome |
| 22EPA096CP6 | PRJNA1231000 | SRR32681753 | Human stool | Genome |
| 22EPA096CP7 | PRJNA1231000 | SRR32681752 | Human stool | Genome |
| 22EPA096CP8 | PRJNA1231000 | SRR32681812 | Human stool | Genome |
| 22EPA096CP9 | PRJNA1231000 | SRR32681811 | Human stool | Genome |
| 22EPA096CPSA1 | PRJNA1231000 | SRR32681810 | Human stool | Genome |
| 22EPA096CPSA10 | PRJNA1231000 | SRR32681771 | Human stool | Genome |
| 22EPA096CPSA6 | PRJNA1231000 | SRR32681765 | Human stool | Genome |
| 22EPA096CPSA7 | PRJNA1231000 | SRR32681770 | Human stool | Genome |
| 22EPA096CPSA8 | PRJNA1231000 | SRR32681768 | Human stool | Genome |
| 22EPA096CPSA9 | PRJNA1231000 | SRR32681767 | Human stool | Genome |
| 23EPA116CP01 | PRJNA1231000 | SRR32681766 | Human stool | Genome |
| 23EPA116CP02 | PRJNA1231000 | SRR32681797 | Human stool | Genome |
| 23EPA116CP03 | PRJNA1231000 | SRR32681796 | Human stool | Genome |
| 23EPA116CP04 | PRJNA1231000 | SRR32681795 | Human stool | Genome |
| 23EPA116CP05 | PRJNA1231000 | SRR32681794 | Human stool | Genome |
| 23EPA116CP06 | PRJNA1231000 | SRR32681793 | Human stool | Genome |
| 23EPA116CP07 | PRJNA1231000 | SRR32681792 | Human stool | Genome |
| 23EPA116CP08 | PRJNA1231000 | SRR32681790 | Human stool | Genome |
| 23EPA116CP09 | PRJNA1231000 | SRR32681789 | Human stool | Genome |
| 23EPA116CP10 | PRJNA1231000 | SRR32681788 | Human stool | Genome |
| 23EPA116CP11 | PRJNA1231000 | SRR32681787 | Human stool | Genome |
| 23EPA116CP12 | PRJNA1231000 | SRR32681786 | Human stool | Genome |
| 23EPA116CP13 | PRJNA1231000 | SRR32681785 | Human stool | Genome |
| 23EPA116CP14 | PRJNA1231000 | SRR32681784 | Human stool | Genome |
| 23EPA116CP15 | PRJNA1231000 | SRR32681783 | Human stool | Genome |
| 23EPA116CP17 | PRJNA1231000 | SRR32681782 | Human stool | Genome |

| | | | | |
|---|---|---|---|---|
| 23EPA116CP18 | PRJNA1231000 | SRR32681781 | Human stool | Genome |
| 23EPA116CP19 | PRJNA1231000 | SRR32681779 | Human stool | Genome |
| 23EPA117CP01 | PRJNA1231000 | SRR32681778 | Human stool | Genome |
| 23EPA117CP02 | PRJNA1231000 | SRR32681777 | Human stool | Genome |
| 23EPA117CP03 | PRJNA1231000 | SRR32681776 | Human stool | Genome |
| 23EPA117CP04 | PRJNA1231000 | SRR32681775 | Human stool | Genome |
| 23EPA117CP05 | PRJNA1231000 | SRR32681774 | Human stool | Genome |
| 23EPA117CP06 | PRJNA1231000 | SRR32681773 | Human stool | Genome |
| 23EPA117CPSA01 | PRJNA1231000 | SRR32681772 | Human stool | Genome |
| 23EPA117CPSA02 | PRJNA1231000 | SRR32681764 | Human stool | Genome |
| 23EPA117CPSA03 | PRJNA1231000 | SRR32681763 | Human stool | Genome |
| 23EPA117CPSA04 | PRJNA1231000 | SRR32681762 | Human stool | Genome |
| 23EPA117CPSA05 | PRJNA1231000 | SRR32681761 | Human stool | Genome |
| 23EPA117CPSA06 | PRJNA1231000 | SRR32681760 | Human stool | Genome |
| 23EPA117CPSA07 | PRJNA1231000 | SRR32681759 | Human stool | Genome |
| 23EPA117CPSA08 | PRJNA1231000 | SRR32681757 | Human stool | Genome |
| 23EPA117CPSA09 | PRJNA1231000 | SRR32681756 | Human stool | Genome |
| 22EPA095CP1 | PRJNA1231000 | SRR32681755 | Human stool | Genome |
| 22EPA095CPSA2 | PRJNA1231000 | SRR32682246 | Human stool | Genome |
| 22EPA095CPSA3 | PRJNA1231000 | SRR32682245 | Human stool | Genome |
| 22EPA095CPSA6 | PRJNA1231000 | SRR32682244 | Human stool | Genome |

**Table S2.** Family-level bacterial population composition and read count in stool microbiomes of three patients and over time in patient-1.

| | No. of reads | | | | |
|---|---|---|---|---|---|
| **Family** | **Patient-1 T1** | **Patient-1 T2** | **Patient-1 T3** | **Patient-2 T1** | **Patient-3 T1** |
| *Actinomycetaceae* | 983 | 64 | - | 47346 | - |
| *Aeromonadaceae* | - | - | - | 2370 | - |
| *Atopobiaceae* | 2658 | - | - | 34328 | - |
| *Bacillaceae* | 55 | - | - | 47475 | - |
| *Bacteroidaceae* | 488596 | 106672637 | 55545179 | 5.1E+07 | 2.9E+07 |
| *Bifidobacteriaceae* | 4128 | 1903623 | 97347 | - | 1956 |
| *Campylobacteraceae* | 37026 | - | - | 34175 | 26894 |
| *Candidatus Nanosynbacteraceae* | - | - | - | 5487 | - |
| *Candidatus Saccharibacteria* | 1241 | 296 | - | 18372 | - |
| *Carnobacteriaceae* | 4012 | - | - | 2092 | - |
| *Clostridiaceae* | 1576 | 752806 | 8104 | 21267 | 73136 |
| *Clostridiales_Family_XIII_Incertae_Sedis* | - | - | - | 21496 | - |
| *Coriobacteriaceae* | - | 1598 | 289 | - | 17377 |
| *Desulfovibrionaceae* | - | - | 268358 | - | 334143 |
| *Eggerthellaceae* | - | - | - | 766 | 2103 |
| *Enterobacteriaceae* | 7.1E+07 | 406514 | 3253933 | 1987871 | 5164904 |
| *Enterococcaceae* | - | 1250 | - | 7028 | - |
| *Erysipelotrichaceae* | - | 104187 | 23014 | 220416 | 38455 |

| | | | | | |
|---|---|---|---|---|---|
| *Firmicutes_unclassified* | - | - | - | - | 10852 |
| *Fusobacteriaceae* | 456656 | 66160 | - | 15039 | - |
| *Lachnospiraceae* | 156207 | 2869579 | 311630 | 15934 | 173823 |
| *Lactobacillaceae* | - | 23898 | - | - | - |
| *Leptotrichiaceae* | - | - | - | 76759 | - |
| *Micrococcaceae* | 299 | 10045 | - | 1871 | - |
| *Microcoleaceae* | - | - | 733 | - | - |
| *Morganellaceae* | - | 111315 | - | - | - |
| *Neisseriaceae* | - | - | - | 642618 | - |
| *Oscillospiraceae* | - | - | 266070 | - | - |
| *Pasteurellaceae* | 30025 | 3416 | 403 | 2442963 | 22617 |
| *Peptoniphilaceae* | 684 | - | - | 10953 | 7006 |
| *Peptostreptococcaceae* | 22093 | 7459 | 227 | 32902 | 12960 |
| *Porphyromonadaceae* | - | - | - | 675 | - |
| *Prevotellaceae* | 4201 | 1405 | - | 6993 | 384405 |
| *Rikenellaceae* | - | - | - | - | 36275 |
| *Ruminococcaceae* | - | 361725 | 2536493 | - | 133831 |
| *Streptococcaceae* | 82447 | 349149 | 19214 | 219662 | - |
| *Sutterellaceae* | - | 1107158 | 283733 | - | 98533 |
| *Tannerellaceae* | 2920 | 3440379 | 529465 | - | 1600689 |
| *Veillonellaceae* | 2439689 | 3483188 | 723702 | 43487 | 231991 |

**Table S3.** Pathogen detection, description and parameters used in comparisons for stool samples of three patients at infection onset.

| Parameter | Patient-1 T1 | Patient-2 T1 | Patient-3 T1 |
|---|---|---|---|
| *Campylobacter* reads (% of overall reads) | 37,026 (0.05%) | 34,175 (0.06%) | 26,894 (0.05%) |
| *Salmonella* reads (% of overall reads) | 7,637 (0.01%) | No detected | 43,452 (0.08%) |
| % *Campylobacter* MDG completeness (species identified) | 46.3% (*C. jejuni*) | 2.20% | 2.90% |
| *Salmonella* MDG completeness (species identified) | Not recovered | Not recovered | 4.8% (*S. enterica*) |
| *Escherichia* reads (% of overall reads) | 71,126,933 (95%) | - | 4,869,646 (13%) |
| % *Escherichia* MDG completeness (species identified) | 97.1% (*E. coli*) | - | 10.5% (*E. coli*) |
| qPCR Ct value (*Campylobacter*) | 30 | 36 | 40 |
| qPCR Ct value (*Salmonella*) | 31 | 35 | 30 |
| No. of *Campylobacter* isolates | 4 | 18 | 6 |
| No. of *Salmonella* isolates | 3 | 0 | 9 |
| Bristol scale stool score | 7 | 7 | 7 |

**MDG:** metagenome-derived genome; **qPCR:** quantitative polymerase chain reaction

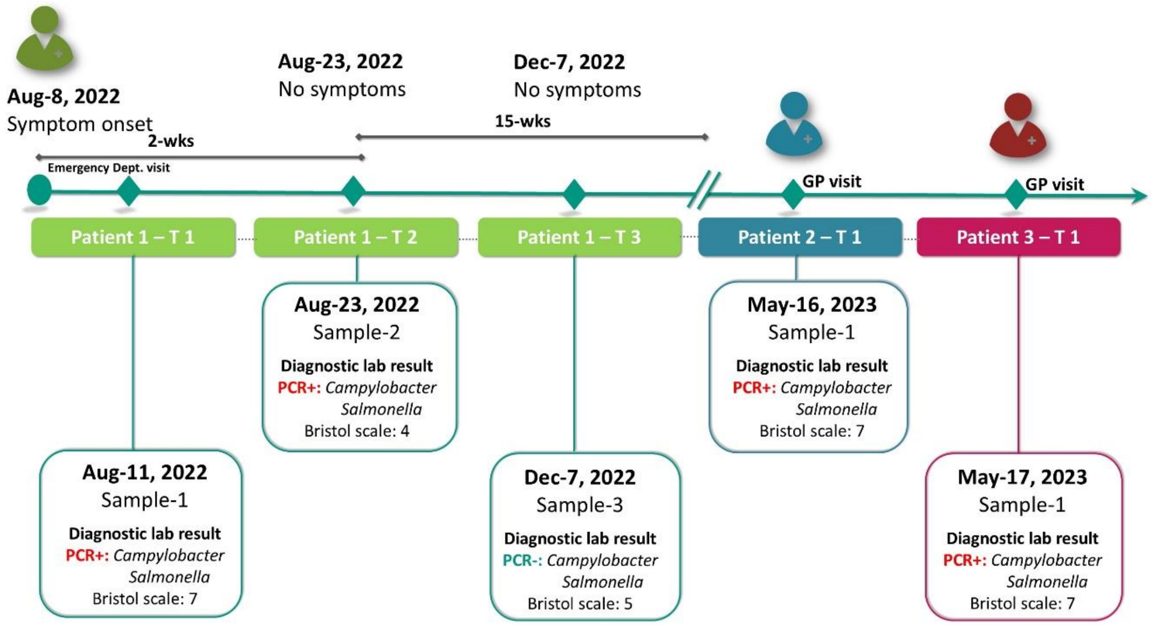


**Figure S1.** Graphical summary of sample collection, patient gastrointestinal illness symptoms and diagnostic laboratory PCR results for three patients with a *Campylobacter* and *Salmonella* gastroenteritis co-infection.

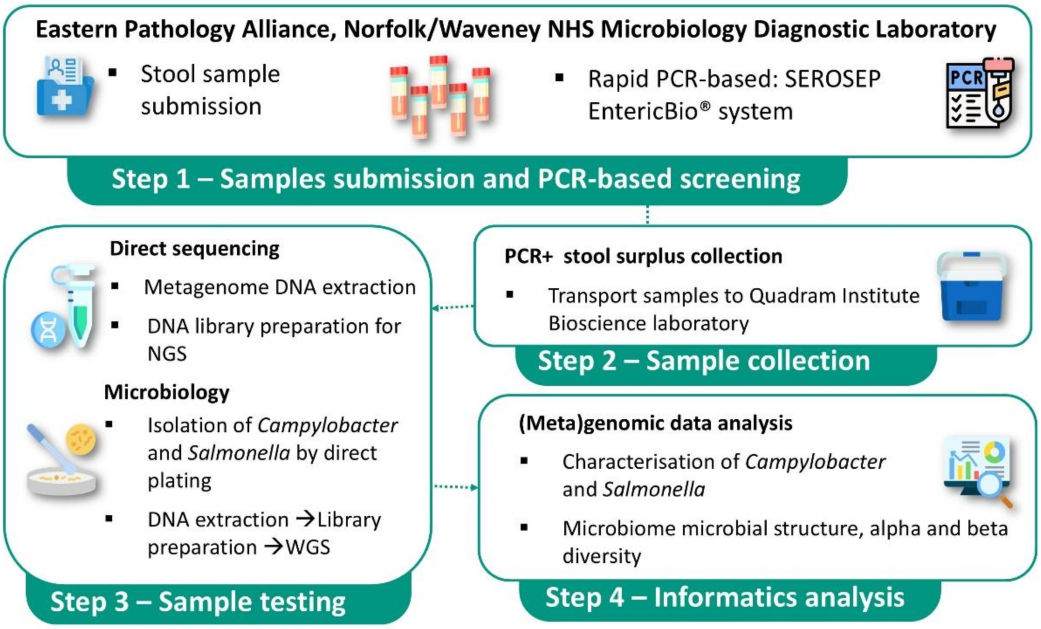


**Figure S2.** Workflow overview of stool sample processing through four general steps from sample collection to informatic analysis.

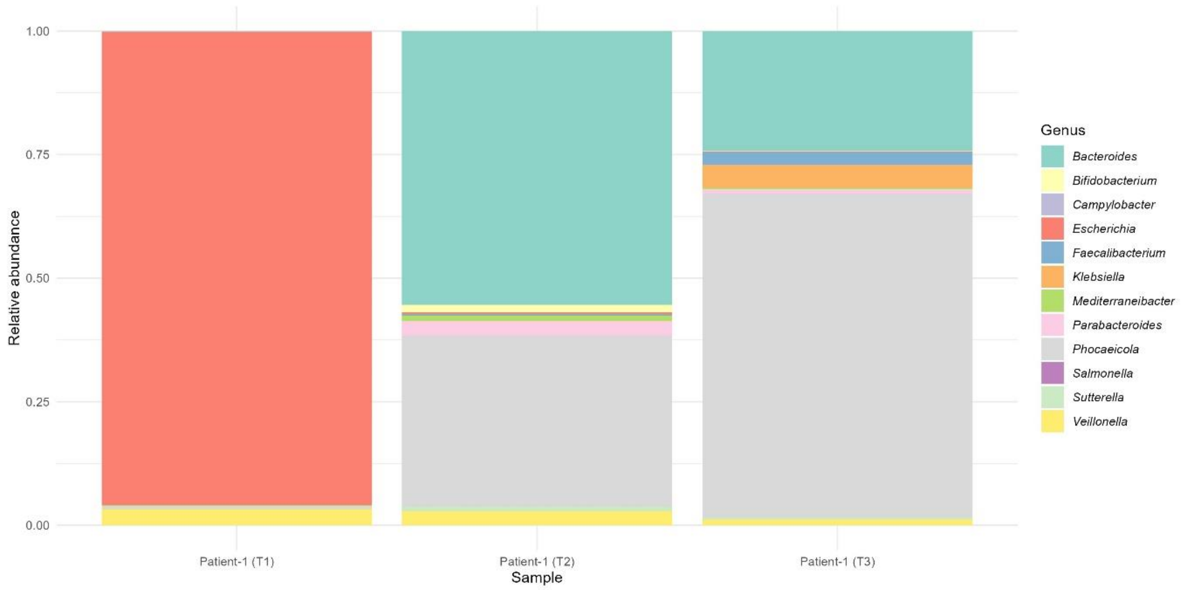


**Figure S3.** Microbiome relative abundance structure in patient-1 at three timepoints (T1, T2, T3) during and after a *Campylobacter* and *Salmonella* co-infection. Relative abundance profiles are shown at the genus level, highlighting the top 10 most abundant genus plus *Campylobacter* and *Salmonella*.

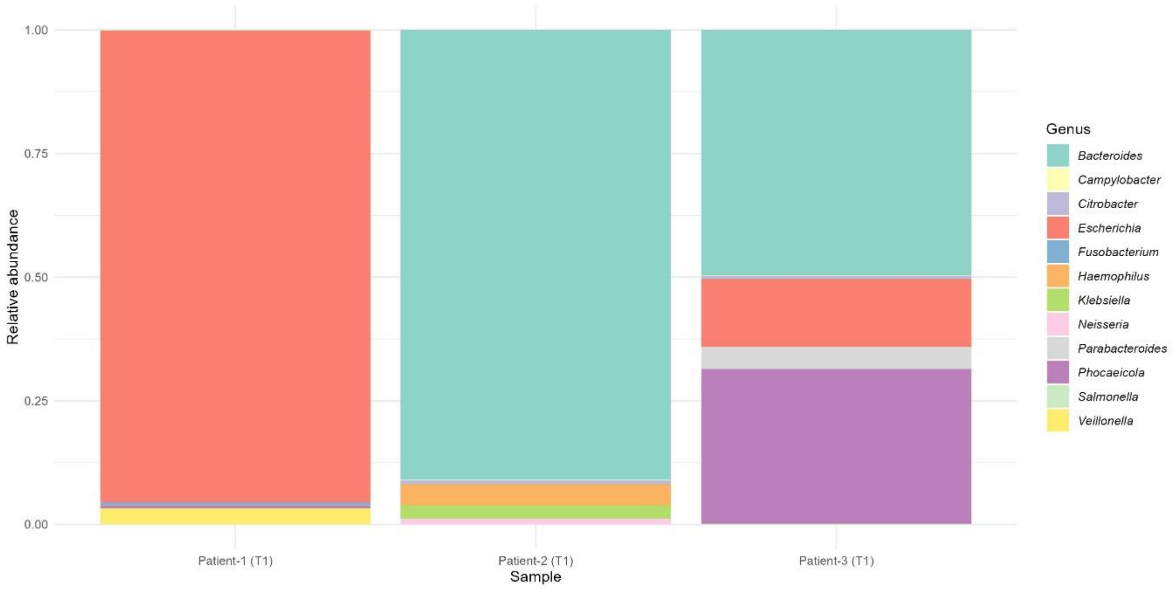


**Figure S4.** Relative abundance microbiome structure of stool samples for three patients' co-infection with *Campylobacter* and *Salmonella*. Relative abundance profiles are shown at the genus level and genus level, highlighting the top 10 most abundant genus plus *Campylobacter* and *Salmonella*.
